# Transcriptional signatures of heroin intake and seeking throughout the brain reward circuit

**DOI:** 10.1101/2023.01.11.523688

**Authors:** Caleb J Browne, Rita Futamura, Angélica Minier-Toribio, Emily M Hicks, Aarthi Ramakrishnan, Freddyson Martínez-Rivera, Molly Estill, Arthur Godino, Eric M Parise, Angélica Torres-Berrío, Ashley M Cunningham, Peter J Hamilton, Deena M Walker, Laura M. Huckins, Yasmin L Hurd, Li Shen, Eric J Nestler

## Abstract

Opioid use disorder (OUD) looms as one of the most severe medical crises currently facing society. More effective therapeutics for OUD requires in-depth understanding of molecular changes supporting drug-taking and relapse. Recent efforts have helped advance these aims, but studies have been limited in number and scope. Here, we develop a brain reward circuit-wide atlas of opioid-induced transcriptional regulation by combining RNA sequencing (RNAseq) and heroin self-administration in male mice modeling multiple OUD-relevant conditions: acute heroin exposure, chronic heroin intake, context-induced drug-seeking following prolonged abstinence, and heroin-primed drug-seeking (i.e., “relapse”). Bioinformatics analysis of this rich dataset identified numerous patterns of molecular changes, transcriptional regulation, brain-region-specific involvement in various aspects of OUD, and both region-specific and pan-circuit biological domains affected by heroin. Integrating RNAseq data with behavioral outcomes using factor analysis to generate an “addiction index” uncovered novel roles for particular brain regions in promoting addiction-relevant behavior, and implicated multi-regional changes in affected genes and biological processes. Comparisons with RNAseq and genome-wide association studies from humans with OUD reveal convergent molecular regulation that are implicated in drug-taking and relapse, and point to novel gene candidates with high therapeutic potential for OUD. These results outline broad molecular reprogramming that may directly promote the development and maintenance of OUD, and provide a foundational resource to the field for future research into OUD mechanisms and treatment strategies.

## Introduction

Opioid use disorder (OUD) currently stands as one of the most significant public health crises facing North America. Over 70,000 people die each year in the USA alone to opioid-related overdose (*1*), and this mortality rate continues to rise, accelerated by the COVID-19 pandemic(*2*). While tragic, overdose deaths represent the tip of the iceberg; for each person lost to opioid overdose, hundreds more are struggling with opioid misuse, and thousands more are receiving opioids for pain management. Unfortunately, treatment for OUD is limited, and sustained abstinence is nearly impossible (*3, 4*). To weather the growing opioid crisis, we desperately need new therapeutic approaches based on fundamental biological insights into how OUD develops and is maintained.

The primary barrier to treating OUD, as with other addictive disorders, is relapse (*5*). Patients struggling with OUD become trapped in a cycle of drug-taking and drug-seeking which is fueled by craving during periods of abstinence. Craving can not only drive ongoing drug use but precipitate relapse even years after cessation. Drug use causes neuroplastic changes in the brain’s reward circuitry that promote craving by tuning motivation towards obtaining drugs of abuse and away from natural rewards and healthy goals (*6, 7*). These drug-induced maladaptive changes are thought to be supported in part by persistent molecular adaptations within cells that comprise the reward circuitry (*8*). Numerous molecular mechanisms that drive reward circuit dysfunction in addiction have been identified for psychostimulants, but comparably much less is known for opioids (*9*). The majority of our knowledge of molecular changes in animal models of OUD comes from non-contingent exposure conditions and single-brain-region studies. Additionally, few studies have focused on identifying molecular changes that drive drug-seeking and relapse following protracted abstinence.

The goal of the present work was to identify transcriptional mechanisms supporting vulnerability to long-term opioid intake and relapse using a mouse model of OUD. To achieve this, we performed genome-wide transcriptional profiling by RNA sequencing (RNAseq) across six brain regions crucial for reward processing in multiple experimental conditions modeling distinct stages of OUD. Using bioinformatic approaches we investigate transcriptional regulatory processes governing aspects of OUD, and link behavioral profiles reflective of OUD with gene expression patterns. We then overlay these findings in mice to published RNAseq and genome-wide association studies (GWASs) of human OUD and establish that the mouse models successfully recapitulate important swaths of the molecular pathology of the human syndrome, and identify highly relevant interventional targets. Together, this study provides fundamentally new insight into the molecular underpinnings of OUD and opens several novel avenues for therapeutic development.

## METHODS

### Animals

Adult male C57BL/6J mice (9-10 weeks old at the beginning of experimentation) were housed on a 12 h reverse light-dark cycle (lights on at 19:00) and maintained on *ad libitum* food and water access, except during pre-training wherein mice were food restricted as described. Mice were group housed until catheterization, after which point they were single housed for the duration of the study.

### Heroin Self-Administration Procedure

#### Pre-training

Mice were maintained on mild, overnight food restriction and first trained to lever press for a food reward in operant boxes (Med Associates; Fairfax, VT). Two levers were presented to mice at the beginning of the session, and responding on the active lever led to the delivery of a chocolate pellet (Bio-Serv; Flemington, NJ) according to a fixed-ratio 1 (FR1) schedule of reinforcement. Responding on the inactive lever had no consequence. Sessions lasted for 1 h or until 30 rewards had been earned. Mice were trained until acquisition criteria for lever pressing had been met (30 pellets earned in two consecutive sessions; 3-4 sessions on average required). Following acquisition, mice were returned to *ad libitum* food access for the remainder of the study.

#### Jugular vein catheterization

Following acquisition of responding for food mice, mice (N=72) maintained under inhaled isoflurane anesthesia (2%) received surgery to implant a pre-constructed intravenous catheter (Strategic Applications Incorporated; Lake Villa, IL; SBD-05C) in the right jugular vein. Catheter tubing (0.013” inner diameter) was threaded subcutaneously over the right shoulder and inserted 1 cm into the jugular vein. The catheter cannula exited the skin from the animal’s mid-back. Ketoprofen (5 mg/kg) was injected subcutaneously upon completion of surgical procedures as post-operative analgesia. Following surgery, mice were single-housed for the duration of the study and given 3-4 days of recovery. During recovery, catheters were flushed once daily with 0.03 ml heparinized saline (30 U) containing ampicillin antibiotic (5 mg/ml) to forestall infection.

#### Heroin self-administration

In 15 daily 4 h sessions, responding on the active lever, which previously delivered food, now delivered an intravenous infusion of heroin (0.05 mg/kg/infusion, 24.5 μl infusion volume) on a FR1 schedule of reinforcement. Infusions were signaled by the illumination of a cue light located above the active lever and were followed by a 20 s timeout, during which the cue light remained illuminated and levers remained extended. During timeout, responses on the active lever were recorded but had no consequence. Throughout testing, responses on the inactive lever were recorded but had no consequence. Catheters were flushed with heparinized saline (30 U) before and after self-administration sessions. For the first two sessions, the maximum number of infusions was capped at 60 to minimize overdosing. Thereafter, maximum infusions were capped at 100.

Following the 15^th^ self-administration session, saline and heroin self-administering animals were separated into two withdrawal conditions: 24 h withdrawal (S24 or H24) or extended 30 day homecage withdrawal (i.e., forced abstinence). In the extended withdrawal condition, 30 days after the last self-administration session, mice were further subdivided into two groups receiving a challenge injection of either saline (SS or HS) or heroin subcutaneously at a dose of 1 mg/kg (SH, HH) immediately followed by a drug-seeking test. Mice in all 4 groups were balanced for baseline heroin intake. Here, mice were placed back into their original self-administration chambers with levers extended for 2 h. During this test, lever pressing is recorded but does not lead to drug delivery or cue light illumination, thus measuring anticipatory responses for heroin. Animals were run in two cohorts to ensure similar ages at time of euthanasia between the 24 h and 30 d withdrawal groups; while the first cohort was undergoing 30 d withdrawal, the second cohort was being tested. 21 mice were removed from the study due to catheter patency loss, poor behavioral outcomes, or death unrelated to behavioral procedures.

### RNAseq and Differential Expression Analysis

Immediately after the drug-seeking test, mice were removed from operant boxes and killed via cervical dislocation. Brains were rapidly extracted on ice, sectioned into 1 mm thick coronal slices using a brain matrix, and brain punches of regions of interest were collected and flash-frozen on dry ice. Unilateral midline punches were taken from medial prefrontal cortex (mPFC; 12 gauge), and bilateral punches were taken from nucleus accumbens (NAc; 14 g), dorsal striatum (dStri; 14 g), basolateral amygdala (BLA; 15 g), ventral hippocampus (vHPC; 16 g), and ventral tegmental area (VTA; 16 g). One BLA sample was lost during tissue collection.

Total RNA was isolated and purified using an RNeasy Micro Kit (Qiagen). All RNA samples used for RNAseq had 260/280 values >1.6 (Nanodrop), and RIN values >8.1 (Tapestation). Library preparation and sequencing was performed by Genewiz/Azenta (Chelmsford, MA). Using a minimum of 200 ng RNA, sequencing libraries were generated for each sample individually with an Illumina Truseq Gold kit with ribosomal RNA depletion. Sequencing was conducted on an Illumina Hi-seq machine with a 2×150 bp paired-end read configuration. All samples (305 in total) were multiplexed and run concurrently to produce 40M paired-end reads per sample. Raw sequencing reads from mice were mapped to mm10 using HISAT2. Counts of reads mapping to genes were obtained using htseq-count against Ensembl v90 annotation. Quality control was performed using FastQC. Four VTA samples did not meet QC criteria or were depleted for dopaminergic gene expression (V-261, V-258, V-279, V-280) and were excluded prior to differential expression analysis. Normalized reads were filtered to include only the top third of the genome, yielding a minimum base mean of ~12, and a total list of ~17,800 genes. Unless otherwise specified, significance for differentially-expressed genes (DEGs) was set at log2FC>30% and p<0.05. Differential expression was performed on this filtered gene list in R version 4.0.2 using the DESeq2 package (*10*), with built-in independent filtering disabled.

### Biotype and Cell-Type Enrichment

For gene biotype analysis, gene lists were filtered across all conditions into 6 biotype groupings according to the Ensembl database: lncRNA (combined antisense, bidirectional promoter lncRNA, sense intronic, and processed transcripts), sncRNA (miRNA, misc RNA, snRNA, and snoRNA), pseudogene (processed pseudogene, unprocessed pseudogene, transcribed processed pseudogene), to be experimentally confirmed (TEC), and protein coding. For cell-type enrichment, DEG lists were merged with a comprehensive single-cell RNAseq database (*11*), which identified between 850-1,050 cell-type-specific genes for astrocytes, endothelial cells, microglia, neurons, oligodendrocytes, and oligodendrocyte precursor cells in brain.

### Gene Ontology Analysis

EnrichR Biological Processes 2021 database and Molecular Functions 2021 was used for gene ontology analyses. Criteria for presentation of GO terms are as described. Specific terms presented throughout results are summarized from output lists containing redundancies.

### Upstream Regulator Analysis

Predicted upstream regulators were identified using Ingenuity Pathway Analysis (Qiagen, Fredrick, MD). Genes included in analyses met a criterion of 30% fold change and p<0.05. Only upstream regulators considered “molecules” (genes and proteins) were examined. Upstream regulators considered for analyses met a Benjamini-Hochberg-corrected p-value of p<0.05 and had a predicted activation score of >0 or inhibition score of <0. Union heatmaps of upstream regulators for each brain region were generated by creating a reference list of all upstream regulators present in each condition, and merging each condition with this reference list.

### Rank-Rank Hypergeometric Overlap

RRHO provides the ability to compare gene expression profiles of two conditions in a threshold-free manner to identify the degree and significance of overlap (*12*). RRHOs were generated to compare transcriptomic overlap across DEG lists for two distinct experimental conditions or brain regions (e.g., SH x HH compares DEG lists from SHvsSS against DEG list from HHvsSS). RRHO plots were generated using the RRHO2 package using standard settings (github.com/RRHO2/RRHO2).

### Generation of the Addiction Index

The R package ‘psych’ was utilized to conduct factor analysis to reduce the dimensions of the behavioral variables in the dataset. All animals with the exception of one outlier (2-31) were included in the analysis. Behavioral measures for IVSA included were: Total infusions, Total intake, Total active lever presses, Total inactive lever presses, Total timeout responses, Active vs. Inactive lever presses. For Total intake, an additional variable referred to as Intake-or-not was included to denote whether the total intake was greater than 0. Behavioral measures included from drug-seeking tests were: Previously active lever presses, inactive lever presses, and Previously Active vs. Inactive lever presses. To determine the optimal number of factors, the function ‘vss’ (Very Simple Structure) of the package ‘psych’ was utilized. The number of factors was set to 3. To determine the factors, the function ‘fa’ was run with the factoring method set to ‘minchi’. Factor scores across all 3 factors for individual animals were estimated using the ‘Bartlett’ method. To compute the addiction index (AI), factor scores were first transformed to eliminate negative values, which resulted in values ranging from 0 to 1 for each factor. The product of the transformed factors was calculated for each animal to obtain the AI. The Voom Limma package in R was used to determine the genes associated with factors. The factors 1, 2 and 3 along with the AI were included as a covariate to the Voom Limma regression model to obtain genes associated with the factors and AI.

### Pattern Analysis: Alluvial Plots

To generate alluvial plots, genes were categorized for significance (p<0.05) in each experimental condition and patterns of common regulation across conditions were mapped using ‘alluvial’ package in R (*13*).

### Comparisons with Human RNAseq OUD Datasets

Human RNAseq data from the PFC and NAc were accessed from (*14*). To compare heroin IVSA gene expression associations with human opioid use disorder (OUD) gene expression associations, genes identified as being significantly up- or down-regulated at p<0.05 were merged. Union heatmaps were generated across human, mouse H24, and mouse HH conditions to identify convergence or divergence across these conditions. Pearson correlations were calculated to quantify strength of correlation between transcriptional signatures.

### Partitioned Heritability Linkage Disequilibrium Score Regression

Partitioned heritability Linkage Disequilibrium Score regression (LDSC) is a modified version of LDSC regression that partitions the heritability of a trait by a set of annotated single nucleotide polymorphisms (SNPs) (*15*). It then tests for enrichment of the annotated SNPs in the heritability of the trait using GWAS summary statistics. We defined gene sets of genes positively and negatively associated with addiction index and annotated SNPs (HapMap3 MAF>5% SNPs; (*16*)) within 100kb of the transcribed region of each gene to derive annotation-specific LD scores using 1000 Genomes EUR ancestry as reference (*17*). We also derived background annotations-specific LD Scores for all SNPs and for all genes captured in our RNAseq datasets across brain regions. We then partition heritability of 6 substance use traits, 3 substance use disorder traits, 7 psychiatric traits, and 3 non-psych or substance use traits using publicly available GWAS summary statistics with partitioned LD Score regression conditioning on background LD Scores. For negative control traits, we used left handedness, bone mineral density, and type 2 diabetes (T2D) as non-psychiatric, non-substance use, brain-related (left handedness), and non-brain-related traits (bone density, T2D). Enrichment was calculated as proportion of heritability normalized by proportion of SNPs annotated and p-values were derived from significance testing procedures as implemented in ldsc software and previously described (*15*).

### Transcriptomic Imputation of Opioid Use Disorder GWAS

Transcriptomic Imputation methods use previously calculated associations between genetic variation and gene expression data to impute genetically-regulated gene expression (GREx) given an independent dataset of genotype array or genome sequencing data. Summary imputation methods translate GWAS associations for a trait from the SNP-level to tissue-specific gene expression-level associations using summary statistics. We used Summary-PrediXcan (S-PrediXcan; (*18*)) to impute gene-level associations of human opioid use disorder risk (*19*) from a GWAS comparing opioid dependence cases to opioid-exposed controls. We used brain region-specific elastic net predictor models of 13 brain regions and whole blood from GTEx data (*18, 20, 21*) to compute OUD-GREx associations for each brain region. Z-scores for OUD-GREx were compared to the AI association Z-scores to identify overlapping gene expression signatures.

## RESULTS

### Heroin self-administration

To model OUD, we used an intravenous self-administration (IVSA) paradigm in mice (Fig. 1A). Mice were first briefly trained to perform a lever press for food reward, and then received jugular vein catheterization. After recovery, mice were returned to operant boxes where they could now respond for IVSA of heroin (0.05 mg/kg/infusion; n=27) or saline (n=24) in 15 daily 4 h sessions. Figure 1B demonstrates that heroin IVSA mice showed goal-directed lever pressing unlike saline mice, with more infusions earned (GroupxSession: F_(14,686)_=5.84, p<0.0001; Group: F_(1,49)_=7.85, p<0.01) and more selective responding on the active lever compared to the inactive lever (Two-way ANOVAs LeverxSession; Saline: no main effect of Lever, F_(1,46)_=1.57, *ns*; Heroin: LeverxSession, F_(14, 728)_=52.62, p<0.0001).

**Fig 1.**
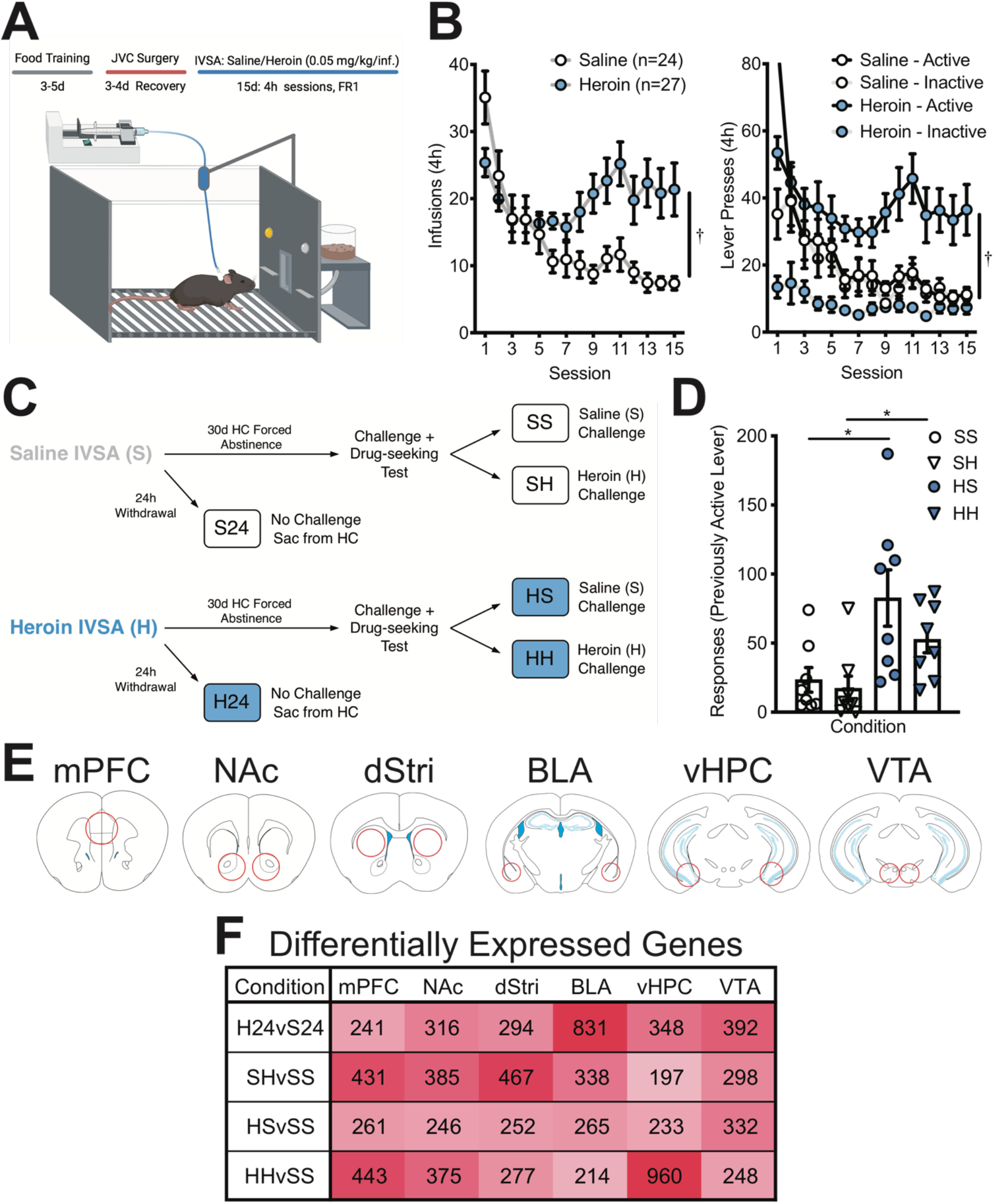
Reward circuitry transcriptome profiling in a mouse model of volitional heroin intake and seeking. (**A**) Intravenous self-administration (IVSA) paradigm. Mice were first briefly trained to lever press for food, then received jugular vein catheterization (JVC) to enable intravenous infusions of saline or heroin contingent upon lever pressing on a fixed-ratio 1 (FR1) schedule of reinforcement. (**B**) Mice performing heroin IVSA took significantly more infusions (left; †p<0.05, Two-way ANOVA) and made significantly more active lever presses compared to inactive lever presses (right; †p<0.05, Two-way ANOVAs) compared to mice performing saline IVSA. (**C**) Schematic describing experimental conditions. After IVSA, mice were split into either 24 h withdrawal (S24, H24) or 30 d homecage (HC) forced abstinence conditions. Mice in the S24 and H24 groups were euthanized 24 h after the final IVSA session. Mice in the 30 d group were further split to receive a challenge injection of either saline (SS, HS) or 1 mg/kg heroin (SH, HH) and were placed back into operant boxes to measure drug-seeking under extinction conditions for 2 h after which they were immediately euthanized. (**D**) Mice that previously self-administered heroin made significantly more responses on the previously active lever compared to saline IVSA mice (*p<0.05, independent samples t-test). (**E**) Six brain regions were extracted for RNAseq: the medial prefrontal cortex (mPFC), nucleus accumbens (NAc), dorsal striatum (dStri), basolateral amygdala (BLA), ventral hippocampus (vHPC), and ventral tegmental area (VTA). (**F**) Summary of the number of differentially expressed genes (DEGs) induced across brain regions and experimental conditions (log2foldchange > 30%, nominal p<0.05).

Following the final IVSA session, mice from both saline and heroin groups were subdivided into short- and long-term homecage forced abstinence conditions of either 24 h or 30 days, respectively (Fig. 1C). Mice in the 24 h withdrawal condition (S24 and H24) were euthanized the following day, directly from the homecage. The 30 d condition was designed to model aspects of incubation of craving for opioids in promoting relapse (*22, 23*). In this condition, mice were further subdivided into two groups, in which they received a challenge injection of either saline (saline-saline, SS, and heroin-saline, HS) or 1 mg/kg heroin (saline-heroin, SH, and heroin-heroin, HH), and were placed back into operant boxes for a 2 h drug-seeking test. During this test, levers were present in the chamber but were both inactive, enabling measurement of anticipatory responses for heroin. Animals with a history of heroin intake showed clear drug-seeking behavior compared to animals that self-administered saline (Fig. 1D; SSvsHS, t(14)=2.66, p<0.05; SHvsHH, t(14), 2.70, p<0.05). This design enabled investigation of the transcriptomic landscape in multiple conditions that model distinct aspects of the OUD syndrome: first-ever heroin exposure (SH), ongoing/early withdrawal from heroin intake (H24), context-induced heroin-seeking following protracted abstinence (HS), and a combination of drug-induced and context-induced heroin-seeking potentially reminiscent of a relapse episode (HH).

### Heroin exposure conditions engage separable gene expression programs in distinct cell-types

Immediately upon completion of the drug-seeking test, animals were euthanized and tissue was dissected for RNAseq (Fig. 1E). Six brain regions that serve as critical hubs in reward-related behavior were extracted and processed for RNAseq: mPFC, NAc, dStri, BLA, vHPC, and VTA. Differential expression analysis (DESeq2) comparing each condition to its respective control identified clear region- and condition-specific changes in transcriptional regulation (Fig. 1F). Across 24 h and 30 d conditions, all brain regions exhibited hundreds of DEGs, demonstrating the ability of heroin intake to cause broad transcriptomic reprogramming throughout the brain’s reward circuitry that persist long after initial intake. We noted that different heroin exposure conditions exerted partly distinct region-specific effects on transcriptional regulation. For example, in the H24 condition, the number of DEGs in the BLA is much higher than in any other brain region studied. Interestingly, first-ever heroin exposure (SH) elicited the weakest DEG response in the vHPC, but heroin re-exposure in animals with a history of heroin intake (HH) caused a strong transcriptional regulation in this brain region, indicating a priming effect of repeated drug exposure on the vHPC transcriptome. In contrast to SH and HH, fewer DEGs are observed for the context-induced drug-seeking (HS) condition. The HS condition is unique compared to SH and HH, in that no drug is on board at the time of tissue harvesting. Thus, these results suggest heroin exposure itself is a major disruptor to transcriptional homeostasis while context-induced drug-seeking is associated with subtler transcriptional changes.

Gene biotype analysis (Fig. S1A) found that the majority (34-88%) of DEGs are protein-coding, but heroin conditions induce particular shifts in biotype induction in a region- and condition-specific manner. The two conditions with highest DEG expression (H24, BLA; HH, vHPC) involved a shift to predominantly protein-coding genes. Additionally, heroin-primed drug-seeking (HH) caused a strong recruitment of non-coding genes in the dStri compared to other exposure conditions. Cell-type enrichment analysis (Fig. S1B) revealed that, in the BLA, the H24 condition uniquely engaged non-neuronal genes compared to other exposure conditions. Additionally, in the mPFC, the SH condition elicited a strong induction of astrocytic genes, but a corresponding depletion of oligodendrocyte genes. In the VTA, we observed an expansion of microglial-enriched genes across exposure conditions peaking with re-exposure to heroin (HH). In the dStri, first-ever exposure to heroin (SH) caused a strong induction of neuronal genes, which contrasts with heroin re-exposure conditions (HH) that exhibited higher astrocytic and oligodendrocytic gene enrichment. Notably, in the NAc, neuronal genes are enriched in the H24 condition, but this contracts for HS and HH conditions suggesting a consolidation of neural transcriptional engagement through abstinence and re-exposure. Overall, these results demonstrate that heroin causes broad transcriptional responses throughout the reward system affecting multiple biological domains, and these effects can shift with chronic intake and re-exposure.

### Heroin intake induces coordinated transcriptional patterning across multiple brain regions

We next asked whether the broad DEG induction observed throughout the reward circuitry involved an ability of heroin to cause coordinated regulation of gene expression across the brain regions examined. Examining the balance of up- and downregulated genes in the six brain regions (Fig. 2A), we noted that the direction of gene expression changes was varied for H24, SH, and HS conditions, but that HH generally suppressed gene expression across all regions. The H24 condition caused exceptionally strong upregulation of genes in the BLA, which contrasts to the generalized downregulation observed across all other conditions in this region. Further, H24 exhibited high concordance of upregulated genes in BLA and vHPC, with 72 genes commonly upregulated in the two regions (Fig. 2B). Additionally, many overlapping BLA and vHPC genes induced at H24 are upregulated across other brain regions (Fig. 2C), including 4 genes (*Hbb-bt, Hba-a1, Depp1*, and *Alas2*) that are also upregulated in 5 of 6 regions studied. First-ever heroin exposure (SH) caused a coordinated transcriptional response across mPFC, NAc, dStri, and BLA, particularly for downregulated genes (Fig. 2B, 2C). Interestingly, this coordinated response was less evident when animals had a history of heroin intake (HH), suggesting a potential weakening of coordinated transcriptional responses as a lasting consequence of repeated heroin intake. However, one gene, *Acer2*, stands out as being consistently suppressed in multiple regions in both SH and HH conditions. *Tmem252* was also suppressed in SH across 5 brain regions, and trending towards significant downregulation in the vHPC (p=0.077). This gene was also implicated as an addiction-relevant target spanning multiple brain regions in our previous paper examining cocaine self-administration (*24*). Despite context-induced drug-seeking (HS) exhibiting few overlapping DEGs, one gene, *Xlr3b* (X-linked lymphocyte-regulated protein 3B), was upregulated across all six brain regions. Although not well characterized, *Xlr3b* is an X-linked gene which may be crucial for neurodevelopment and memory-related cognitive processes (*25*), making it a notable target to regulate context-driven drug-seeking behavior. Enrichr gene ontology (GO) analysis of genes showing multiple overlaps across conditions identified enrichment of genes involved in multiple biological processes related to extracellular matrix (ECM) remodeling, as well as cellular trafficking (chemokines), immune responses (cytokines), and metal homeostasis (Fig. 2D). Together, these results demonstrate that heroin exposure, intake, seeking, and relapse each induce patterns of coordinated gene expression throughout the brain reward circuitry affecting overlapping biological domains.

**Fig 2.**
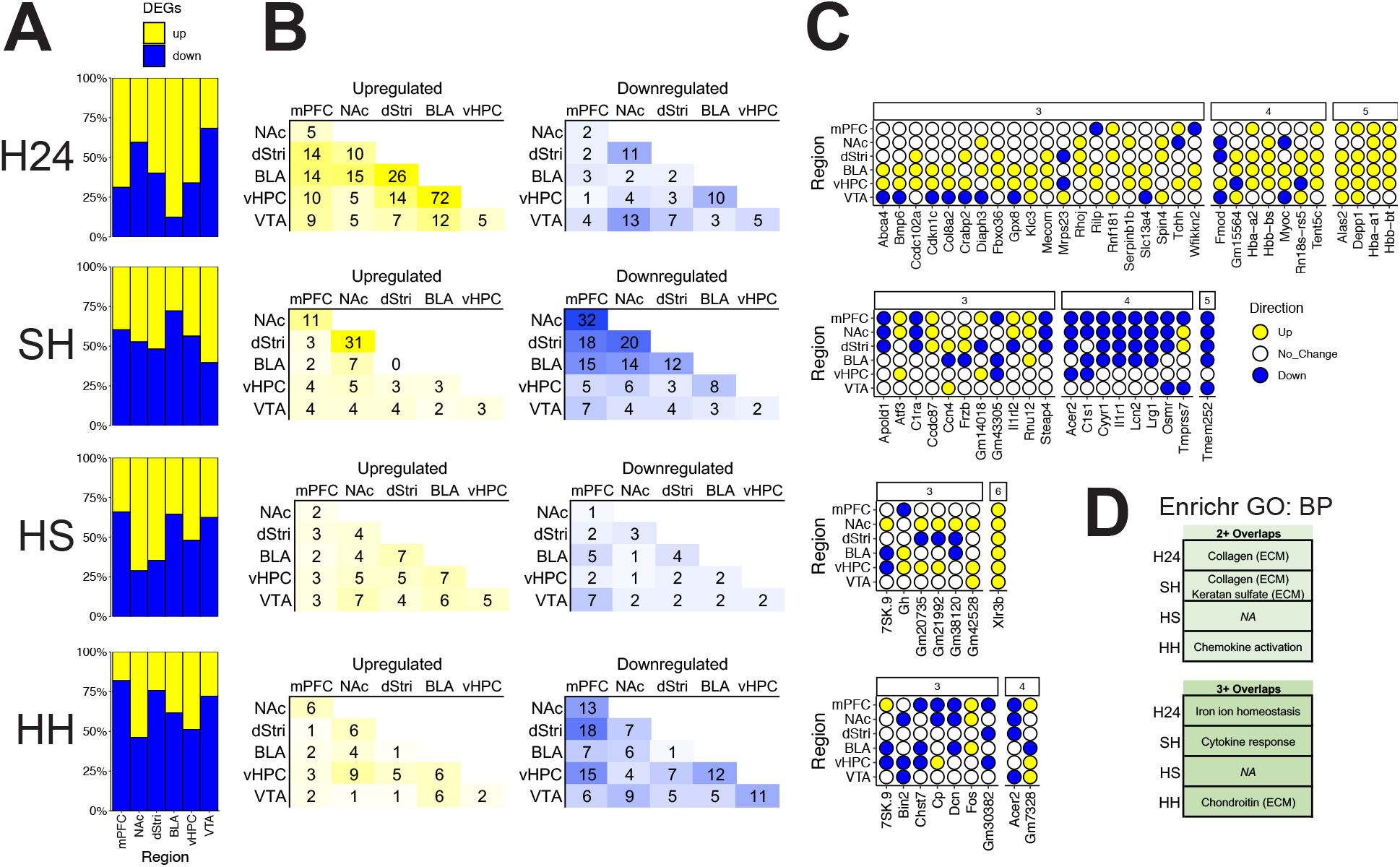
Heroin drives coordinated transcriptional patterning across brain reward regions that shifts with abstinence after a history of intake. (**A**) Proportion of upregulated (yellow) and downregulated (blue) genes across brain regions for each experimental condition. (**B**) Number of shared DEGs identified across paired brain regions as being upregulated (left) or downregulated (right). (**C**) Genes identified as being regulated across 3 or more brain regions, with many showing coordinated expression changes across all regions identified. (**D**) Enrichr gene ontology analysis reveals that genes expressed across multiple brain regions enrich for biological processes related to various extracellular matrix (ECM) processes including collagen, keratan sulfate, and chondroitin, in addition to iron homeostasis and cytokine and chemokine activity.

### Early withdrawal from heroin intake is associated with alterations in ECM biology

We explored how transcriptional signatures varied across each heroin intake and exposure conditions. The H24 condition, reflecting ongoing heroin intake with short-term withdrawal, was considered first based on the distinct control group used. Generally, H24 elicited a unique transcriptional response in comparison to the three forced abstinence conditions (SH, HS, HH; Fig. 3A). This is likely due to the contrasting experience of the groups (and their respective controls) immediately prior to euthanasia: animals in the 24 h condition were killed from their homecage, while animals in the forced abstinence groups experienced an injection and 2 h of behavioral testing. These two experiences may initiate distinct activity/arousal-based transcriptional patterns unrelated to heroin exposure per se. Nonetheless, we noted that certain patterns of gene expression in the H24 condition were either recapitulated or inversed in the forced abstinence conditions. Particularly evident was transcriptional patterning in the BLA and vHPC, where the most upregulated genes in short-term withdrawal (H24) are suppressed by first-ever exposure to heroin (SH) in both the BLA and vHPC. In the vHPC, these genes reverted to being strongly upregulated upon re-exposure to heroin in the HH condition.

**Fig 3.**
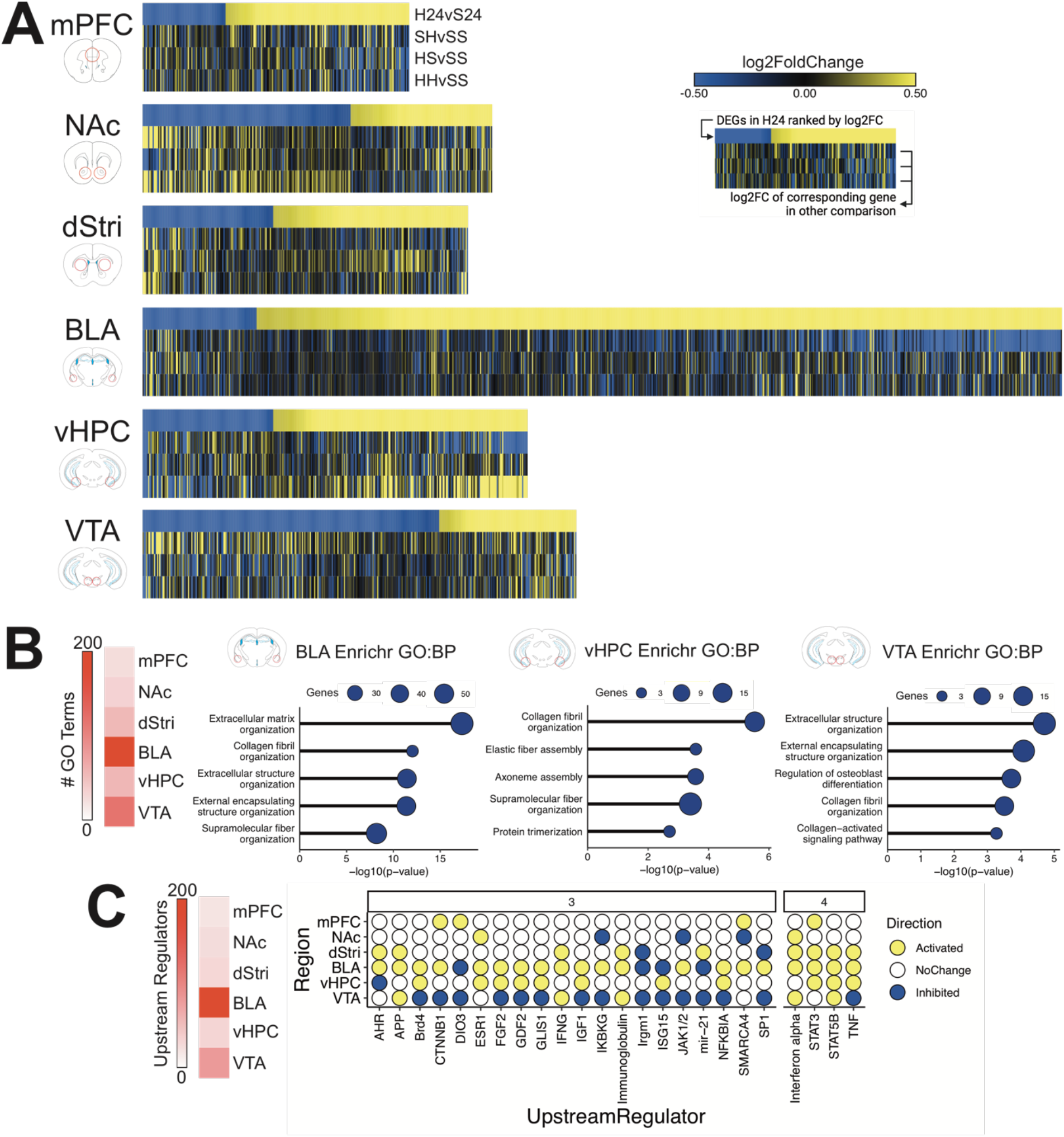
Heroin intake with short-term withdrawal strongly activates transcriptional programs in the BLA and vHPC associated with extracellular matrix (ECM) processes and inflammatory cytokines. (**A**) Heatmaps with top row displaying all DEGs in the H24 condition ranked from lowest to highest log2FoldChange, and log2 FoldChange of corresponding gene in other experimental conditions in subsequent rows. (**B**) Results from Enrichr gene ontology analysis showing enrichment of H24 DEGs in various biological processes. In the BLA, vHPC, and VTA, top enriched terms relate to ECM functions. (**C**) Upstream regulator analysis revealed numerous molecular regulators of gene expression that are activated (yellow) or suppressed (blue) across multiple brain regions.

Gene ontology analysis across brain regions of the H24 condition identified enrichment of numerous biological processes in the BLA, vHPC, and VTA, which predominantly related to the ECM (Fig. 3B). These results indicate that heroin self-administration initiates a multi-region shift in ECM function, consistent with recent suggestions from human postmortem studies (*14*). Heroin self-administration also caused broad shifts in upstream regulator activation which were identified across multiple brain regions (Fig. 3C). The BLA and VTA showed strongest enrichment of upstream regulators, many of which overlapped between the two regions (63/88). However, many of these upstream regulators showed opposite predicted activation or inhibition between the BLA and VTA, with the majority activated in BLA but inhibited in VTA. Upstream regulators that showed enrichment across 4 brain regions, including TNF alpha, STAT5B, STAT3, and interferon alpha, were largely activated throughout the brain. These upstream regulators are associated with cytokine-mediated transcriptional regulation, suggesting coordinated transcriptional responses throughout the brain in mediating cytokine function. Interestingly, cytokines work in partnership with the ECM to maintain structural components of the extracellular milieu and support neural and glial function (*26, 27*). Thus, these results also point to heroin’s ability to disrupt ECM maintenance and cytokine function as a potential brain-wide driver of OUD.

### Segregated transcriptional remodeling across the reward circuitry supports drug- vs. context-specific influences in relapse

A primary goal for this study was to examine transcriptional mechanisms of relapse following protracted abstinence from heroin intake. Thus, we next focused our analysis on mice that underwent 30 days of forced abstinence from IVSA. Using a threshold-free approach, we first asked how transcriptomic remodeling varied across initial heroin exposure (SH), context-induced drug-seeking (HS), and heroin-primed drug-seeking (HH) as a function of experimental condition. We took the approach that SH and HS conditions contain separable components of the HH condition: SH may capture transcriptional regulation mediated specifically by an acute heroin exposure, and HS captures transcriptional regulation mediated specifically by context-dependent memory-related drug-seeking. Rank-rank hypergeometric overlap (RRHO) plots (Fig. 4A) demonstrate that all brain regions show either concordance or no concordance in their transcriptional response across experimental groups – no anti-correlations were apparent. However, we noted that particular brain regions played unique roles in coordinating transcriptional responses to drug experiences. The mPFC, NAc, and dStri showed strongest transcriptional overlap between SH and HH conditions, suggesting that transcriptional remodeling within these regions may be strongly mediated by direct actions of heroin. These effects may indicate that repeated exposure heroin causes iterative engagement of this transcriptional signature, priming neurobiological changes induced in relapse. On the other hand, the BLA and vHPC showed exceptionally strong coordination across HS and HH conditions, implicating a role for transcriptional remodeling supporting drug-seeking specifically in these brain regions. These findings are consistent with prior literature implicating BLA and vHPC in cue- and context-induced drug-seeking (*28, 29*), and extend this idea to molecular reprogramming in these brain regions.

**Fig 4.**
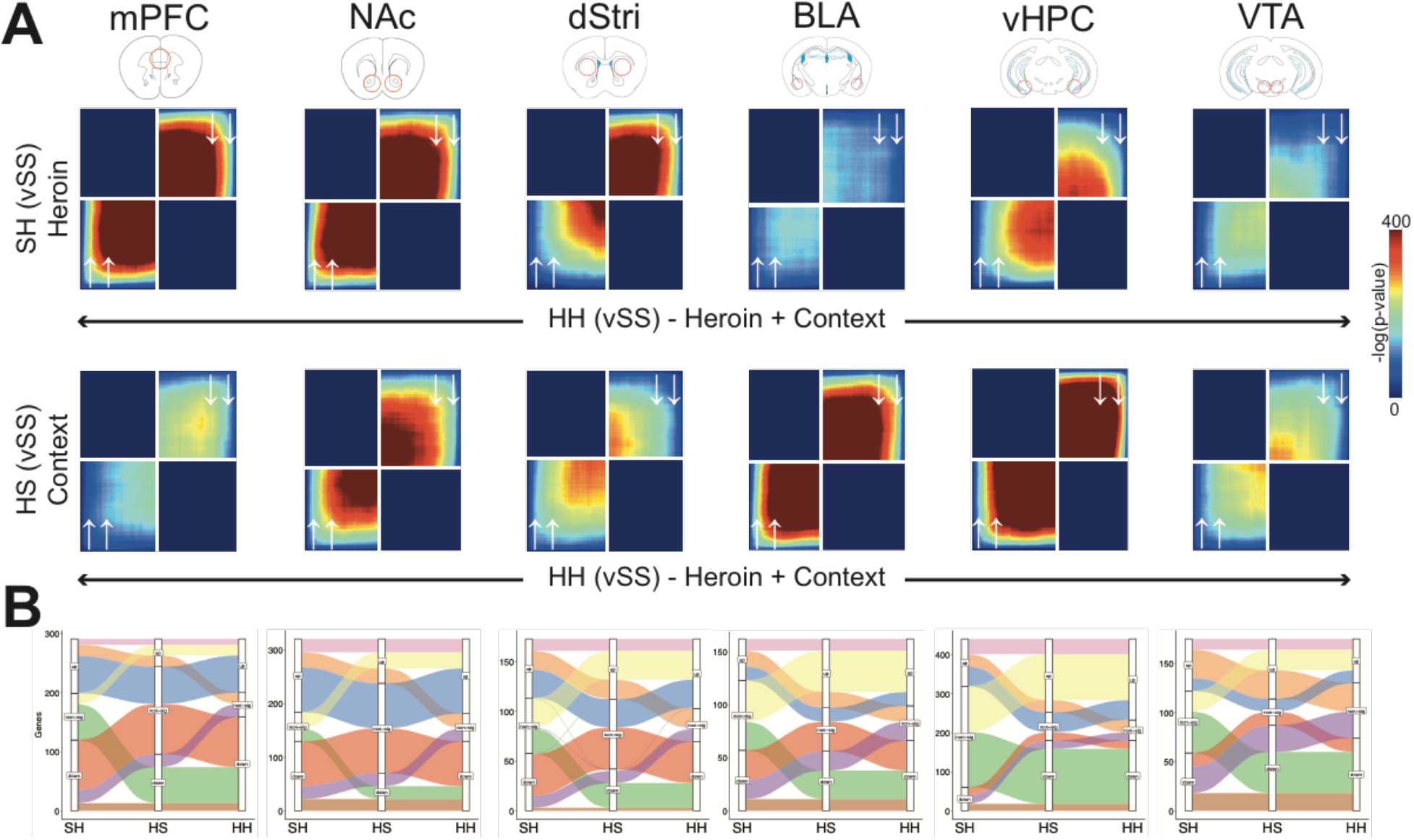
Distinct regional contributions to heroin-priming and context-priming transcriptional regulation associated with relapse. (**A**) Rank-rank hypergeometric overlap plots showing coordinated gene expression between SH and HH conditions (top row) or HS and HH conditions (bottom row) across brain regions. (**B**) Alluvial plots showing clusters of genes (alluvium colors) exhibiting a particular pattern of gene expression through SH, HS, and HH conditions. Graphs are stratified for each experimental condition (x axis) to show the number of genes (y axis) that are upregulated, unchanged, or downregulated. Gene clusters are colored by pattern.

Our experimental design enables profiling of transcriptomic dynamics across what could be considered increasing exposure or severity in OUD: from first-ever experience (SH), to abstinence from chronic intake (HS), and finally to relapse (HH). To examine how sets of genes flow through these heroin experiences, we used a pattern analysis approach to group genes based on their expression pattern of being either significantly upregulated, unchanged, or significantly downregulated across SH, HS, and HH conditions. Alluvial plots in Figure 4B demonstrate that many clusters of genes take unique paths through heroin exposure conditions, and that the magnitude of genes following particular paths shifts across brain regions. All regions have several genes that show consistent and persistent up or downregulation across each heroin exposure condition. Notably very few genes show patterns of inversion of expression, i.e., genes that are up in one condition are typically not down in another condition. Consistent with RRHO analyses, genes in the mPFC, NAc, and dStri that are up- or downregulated in SH are quiescent in HS, but reactivated again in HH upon re-exposure to heroin. Also consistent with RRHO analyses, many genes in the vHPC and BLA that are unaffected by an initial exposure to heroin (SH) exhibit consistent up- or downregulation in HS and HH conditions. Together, these findings suggest that certain groups of genes in the mPFC and NAc, and potentially the dStri, are heavily influenced by the direct effects of heroin which may become primed or desensitized in relapse-like conditions. On the other hand, genes in the BLA and vHPC are primarily induced after prolonged withdrawal from heroin consumption and may relate to context-related memory of drug experience driving drug-seeking.

### Gene priming and desensitization by a distant history of heroin intake

We next examined how heroin exposure and drug-seeking caused shifts in gene priming and desensitization throughout the reward circuitry. Union heatmaps seeded to the HH condition (Fig. S2) showed that gene expression changes across SH, HS, and HH were generally in a similar direction across brain regions. However, hierarchical clustering of these results revealed unique signatures of heroin exposure for each of the brain regions studied (Fig. 4). Strikingly, this approach found that SH and HH conditions generally show major differences in their patterns of gene expression, thus demonstrating how a distant (30 days) prior course of volitional heroin consumption alters the ability of an acute heroin dose to change gene expression. We also noted that context-induced drug-seeking (HS) engaged unique gene sets in multiple brain regions, with the exception of the vHPC, which stood out as having a high degree of overlap with the HH condition. These results are consistent with findings from RRHO analysis between HS and HH conditions, further demonstrating a key role for vHPC as a regulator of relapse-related molecular processes.

Many gene clusters were found to enrich for particular GO processes spanning various domains of biological function (Fig. 5; Supp tables). The most apparent reward circuit-wide changes were in processes related to ECM maintenance, identified as various subdomains in ECM biology including collagen, keratan sulfate, and general changes to the ECM. Gene clusters that were most enriched for DEGs in one particular condition tended to also enrich for certain biological processes. In the mPFC, genes specifically downregulated in HH (cluster 6) enrich for keratan sulfate and sulfur transport, which, by contrast with SH, suggests that repeated heroin exposure primes genes related to these processes. In the dStri, genes predominantly upregulated in SH (cluster 1) enrich for processes related to neurotransmission, which, by contrast with HH, may imply that repeated exposure to heroin diminishes or refines the heroin-induced transcriptional response in this domain. In the VTA, genes downregulated specifically in HS (cluster 3) enrich for collagen-related processes. In the vHPC, clusters of genes that are upregulated specifically in HH enrich for processes related to ECM and vasculature maintenance, while clusters downregulated in HH enrich for processes related to neurotransmission and ion transport. Considering that the ECM is crucial for supporting synaptogenesis, learning, and memory (*30*), the observation of a balanced upregulation of ECM functions with downregulation of synaptic transmission-related processes may imply an ECM-dependent mechanism of neuroplasticity within the vHPC to support relapse. Taken together, these results suggest that heroin engages unique gene priming mechanisms affecting broad neurobiological domains to promote drug-seeking and relapse.

**Fig 5.**
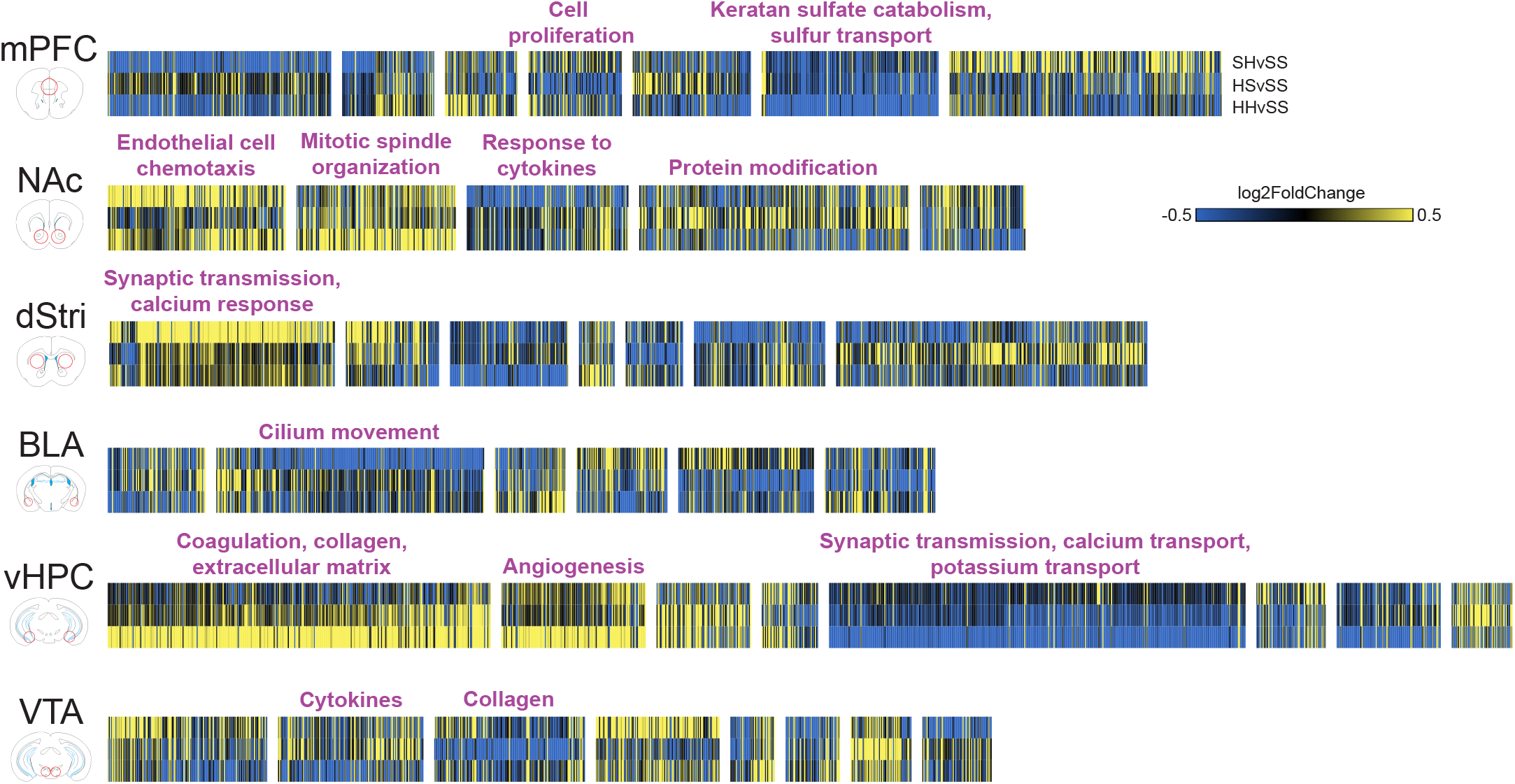
Unique patterns of transcriptional regulation across 30 d withdrawal groups implicate unique biological domains affected by heroin exposure history. Heatmaps show log2FoldChange for genes across SH, HS, and HH conditions. Genes are organized by hierarchical clustering across experimental conditions, with dendrograms cut manually to emphasize clusters. Enrichr GO analysis reveals enrichment of biological processes for particular clusters of genes showing distinct patterns across heroin exposure conditions.

### A history of heroin intake shifts upstream regulator control over transcription in a region-dependent manner

Having identified broad changes to gene expression across heroin exposure conditions, we investigated whether these changes are patterned by particular transcriptional regulatory systems. Using IPA upstream regulator analysis, we deduced whether upstream regulators were significantly activated or inhibited in SH, HS, and HH conditions in the brain regions studied. Many upstream regulators were found to be engaged in a condition-dependent manner across mPFC, NAc, vHPC, and VTA, with fewer total hits for the dStri and BLA (Fig. 6A). In general, the HS condition elicited comparably few upstream regulators across each brain region, with the exception of the VTA. Union heatmaps showing predicted upstream regulators ranked by activation score in HH (Fig. 6A) demonstrate a clear region-specific pattern of activation and inhibition by heroin exposure conditions. Notably, many upstream regulators were unique to the SH condition, again suggesting that the initial transcriptional response to heroin becomes consolidated with repeated exposure. In the mPFC and NAc, strong overlap was observed between SH and HH in upstream regulators predicted to be inhibited or activated. These findings are consistent with transcriptome-wide analyses above demonstrating a heroin-specific response in mPFC and NAc, implicating a critical role for these upstream regulators in mediating drug-priming effects in relapse. On the other hand, the vHPC showed little enrichment of upstream regulators between SH or HS conditions, but strong engagement of many upstream regulators in HH. This contrasts the general patterns observed at the DEG level, where overlap is often observed with the HS condition, indicating that transcriptional control within the vHPC is unique to relapse-like conditions. A different pattern entirely was observed in the VTA, where upstream regulators were enriched across all three conditions, but concordance is most evident for HS and HH.

**Fig 6.**
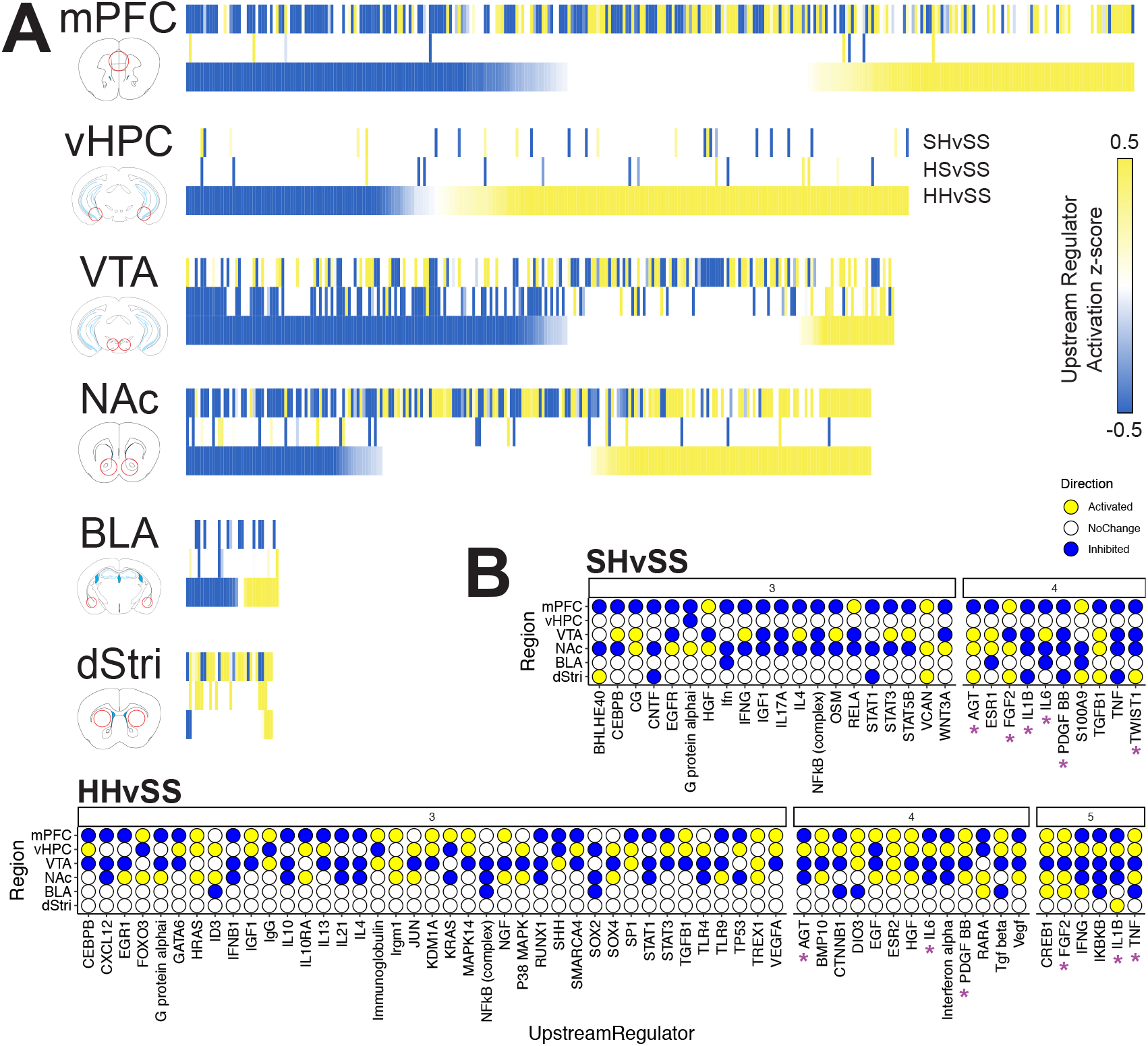
Upstream regulators coordinate distinct components of relapse-related processes in a region-specific manner. (**A**) Heatmaps showing upstream regulators predicted to be inhibited (blue) or activated (yellow) across SH, HS, and HH conditions, ranked by activation Z-score seeded to HH. White indicates upstream regulator not present in experimental condition. (**B**) Upstream regulators identified to exhibit coordinated engagement across 3 or more brain regions in SH and HH conditions (none identified in HS). Stars signify upstream regulators identified to be engaged in ≥4 brain regions in both SH and HH conditions.

We next examined whether heroin exposure conditions drove coordinated changes to upstream regulators. While the HS condition did not elicit significant upstream regulator activation or inhibition across multiple brain regions, numerous instances were observed for SH and HH. Figure 6B shows upstream regulators engaged across 3 or more brain regions for SH and HH, respectively. Interestingly, many upstream regulators were activated or inhibited across up to 4 brain regions in SH and 5 brain regions in HH, suggesting a strong, brain-wide coordination of transcriptional responses to heroin regardless of intake history. We noted that between SH and HH the pattern of activation or inhibition of these multi-region upstream regulators shifts for particular brain regions. For example, the mPFC and NAc shift from mostly inhibited to mostly activated upstream regulators, the vHPC gains numerous upstream regulators, and the dStri loses nearly all upstream regulator engagement. Further, several multi-region upstream regulators were common to both SH and HH, including *Agt, Fgf2, Il1b, Il6, Pdgf-bb*, and *Tnf.* Interestingly, many upstream regulators showing multi-region overlap are commonly inhibited in the mPFC in both SH and HH conditions, implicating common transcriptional responses induced acutely by heroin regardless of a history of heroin intake. It is noteworthy that several transcription factors that have previously been implicated in drug addiction (e.g., CREB1, JUN (an obligatory binding partner for ΔFOSB), and EGR1 are identified in this unbiased dataset as significant upstream regulators of heroin-induced transcriptional regulation (*31*).

### Linking gene expression with addiction-relevant behavioral phenotypes

Thus far, we examined how independent experimental groups reflective of unique OUD conditions influenced transcriptional regulation in brain reward regions. To extend this analysis, we examined how gene expression could predict addiction-relevant behavioral phenotypes and harness individual variability in drug-taking and drug-seeking. Using exploratory factor analysis, we collapsed multidimensional behavioral data collected from animals in all experimental conditions into latent variables (factors) that captured variability across behavioral measures (summarized in Fig. 7A). Three factors captured the most variability in the dataset, and could be considered components of behavioral phenotypes relevant to promoting drug-taking and drug-seeking. Factor 1 was associated with measures related to discrimination of active and inactive lever pressing during self-administration and drug-seeking, and was thus considered a measure of “Selectivity” in responding. Factor 2 was associated with total heroin consumed and was thus considered a measure of “Intake”. Factor 3 was associated with number of responses made, particularly during timeouts between periods of drug availability, and was thus considered a measure of response “Vigor”. To create an individual score for each animal, these factor values were combined into a composite “addiction index” (AI) score ((*24*); Fig. S3A). This AI score scaled across factors, in that higher levels of each factor contribute to a higher AI score and vice versa (Fig. S3B). Importantly, AI scores also scaled with raw behavioral measures, such that higher AI scores were generally associated with higher levels of responding, more lever discrimination, higher timeout responding, and higher drug-seeking (Fig S3C). This approach thus captured aspects of addictionrelevant behavioral phenotypes in a single score for individual animals.

**Fig 7.**
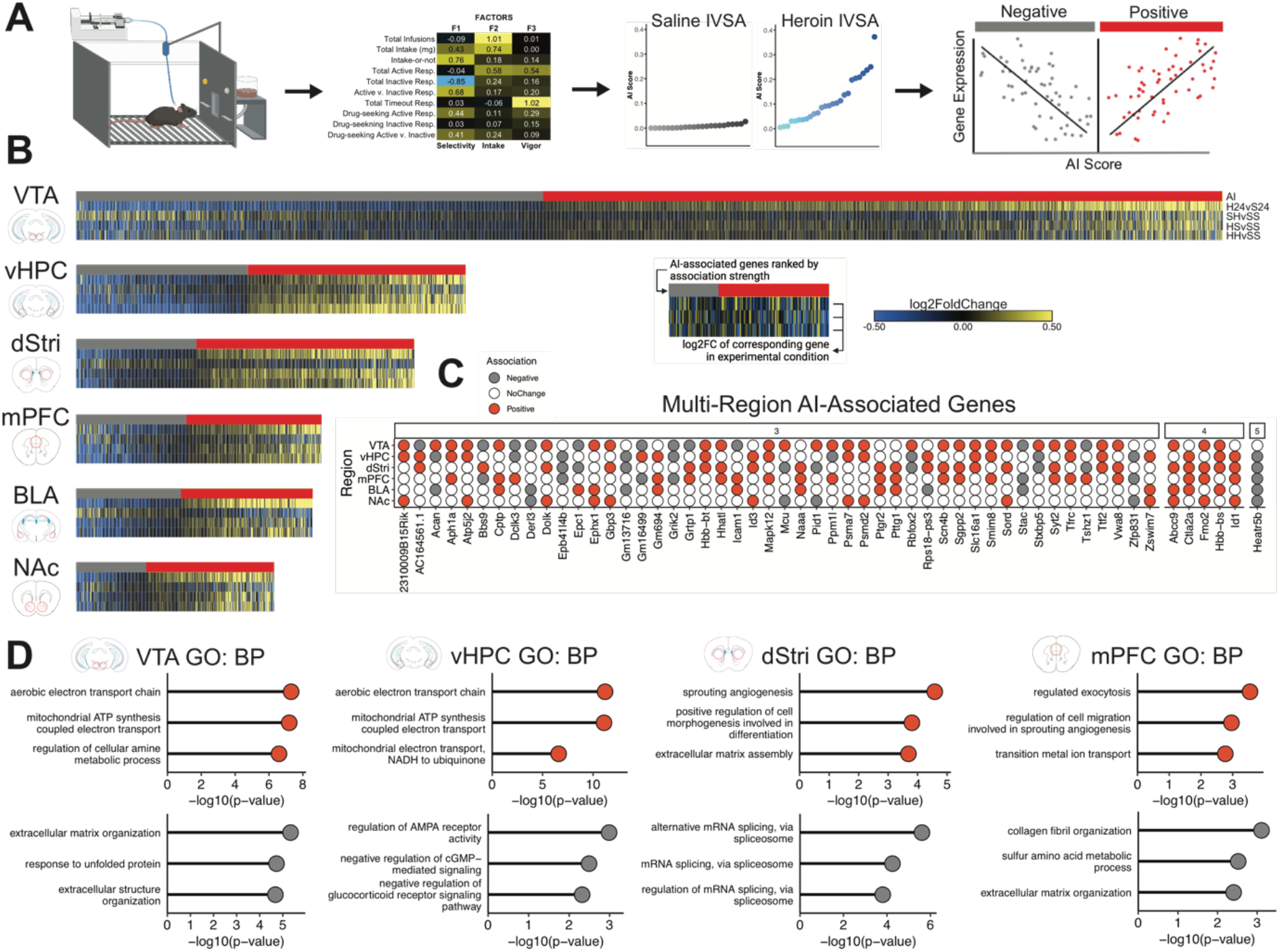
Genes associated with addiction-relevant behavior enrich for processes related to energetic utilization and extracellular matrix function across multiple regions. (**A**) Approach to identify genes associated with addiction-relevant behavioral outcomes of IVSA. Factor analysis was used to collapse all behavioral variables collected during IVSA into three primary latent variables related to “Selectivity” of responding for heroin, total heroin “Intake”, and response “Vigor”. Factors were then combined to generate a single latent variable, termed the “Addiction Index” (AI), for each animal individually, and these scores were higher in animals that selfadministered heroin (middle right, blue symbols, shaded by relative AI score) compared to saline (middle right, gray symbols). Linear regression was then used to identify genes in each brain region that were significantly positively or negatively associated with the AI (slope > 20%, nominal p<0.05). (**B**) Heatmaps showing AI-associated genes (top row; positive in red, negative in gray) ranked by most negative to most positive, and log2FoldChange of corresponding gene from DEG analysis from all experimental conditions (subsequent rows; downregulated in blue, upregulated in yellow) across all brain regions. (**C**) Genes identified to be positively (red) or negatively (gray) associated with the AI across ≥3 brain regions, showing strong coordinated patterns throughout the reward circuit. (**D**) Enrichr GO analysis on positively-associated (red, top graphs) and negatively-associated (gray, bottom graphs) AI genes identifies enrichment of biological processes related to extracellular matrix function and metabolic activity across multiple brain regions.

We then used linear regression to link gene expression with AI scores. Notably, because this approach restricts analysis to genes associated with behavior rather than strictly differential expression relative to a control condition, we were now able to examine transcriptional regulation across all primary experimental conditions (H24, SH, HS, and HH). Heatmaps in Figure 7B show numerous genes identified as being positively associated with AI (red; higher AI = higher expression) or negatively associated with AI (gray; higher AI = lower expression), and their respective expression values across each experimental condition in blue (downregulated vs. control) or yellow (upregulated vs. control). In all drug-taking conditions (H24, HS, and HH), positively associated genes are typically upregulated, while negatively associated genes are typically downregulated. This suggests that heroin intake drives expression of a set of addiction-relevant genes which are maintained as primed or desensitized throughout abstinence and re-engaged upon drug-seeking. Certain regions also showed differing degrees of coordinated activation or inhibition of addiction-related genes in a condition-specific manner. For example, ongoing intake (H24) engages many AI genes in VTA, BLA, and NAc, which contrasts with relapse-like conditions (HH) causing the strongest engagement of AI genes in the vHPC. Acute exposure (SH), on the other hand, causes a largely distinct pattern of AI-associated gene expression compared to those conditions with a history of heroin intake. However, the ability of SH to engage these genes, rather than having no effect on them, is somewhat surprising. This suggests that regardless of a history of heroin intake, initial exposure to heroin engages addiction-relevant gene expression systems throughout the brain reward circuit which become refined and potentially amplified with increasing exposure.

Threshold-free RRHO analysis found that AI-associated genes were largely independent across brain regions, but identified a notable coupling of positive-AI genes between dStri and vHPC, as well as broad concordance between VTA and mPFC (Fig. S4). We identified 54 AI-associated genes that spanned at least 3 brain regions which were mostly consistent in being either positively or negatively correlated with the AI (Fig. 7C). 5 genes (*Hbb-bs, Ctla2a, Id1, Fmo2*, and *Abcc9*) stood out as being positively associated across 4 brain regions, and 1 gene *(Heatr5b*) was negatively associated across 5 regions. *Hbb-bs* was also identified as being upregulated across multiple brain regions in H24 (Fig. 2). As well, we noted more positive (121) compared to negative (48) AI genes when examining overlaps across brain regions, despite the largely balanced number of positive and negative AI genes identified for each region (Fig. 7B). This may imply a reward circuit-wide coordination of gene expression that promotes addiction-like behavior. Enrichr GO analysis of AI-associated genes revealed a strong enrichment of positive-AI genes in biological processes related to energy utilization and mitochondrial function (Fig. 7D). These analyses also identified biological processes related to ECM function enriched in positive-AI genes in the dStri, and negative-AI genes in the mPFC and VTA, further implicating ECM dysfunction in OUD-related processes.

### Integration with human RNAseq and GWAS data reveals convergence with findings in mice as well as key conserved addiction signatures and gene targets for treating OUD

We next integrated our heroin IVSA RNAseq datasets with genome-wide assays from OUD patients to identify commonly regulated genes that may promote addiction (Fig. 8A). First, we compared how gene expression across heroin IVSA conditions compared with human OUD for mPFC and NAc using a recent RNAseq dataset from postmortem brain tissue of OUD patients (Seney, Logan, 2021). Patients in these studies had ongoing OUD and died following opioid overdose. Thus, we determined the H24 and HH conditions, which reflect ongoing heroin intake and re-exposure with heroin onboard, respectively, to be the most relevant comparison conditions to identify convergent patterns of gene expression induced by OUD. Pearson correlation analysis in the mPFC did not identify a correlation between OUD and H24 (R=0.029, *ns*), but a slight negative correlation between OUD and HH (R=-0.054, p<0.01). On the other hand, in the NAc we found a robust positive association between OUD and H24 (R=0.216, p<0.0001) and a negative correlation between OUD and HH (R=-0.199, p<0.0001). We then merged significant DEGs from OUD patients and mouse experimental conditions to generate union heatmaps presented in Figure 8A. Consistent with correlation analysis, we observed a striking overlap of DEGs between human OUD and H24 conditions in the NAc, while the HH condition showed primarily opposite regulation to the human OUD condition. In contrast, less DEG overlap was observed between mouse and human datasets for the mPFC. Using Enrichr GO analysis, we then examined biological process enrichment in genes identified in significantly correlated conditions. Enrichr GO analysis of concordant genes between human and mouse H24 conditions from the NAc identified biological processes related to nervous system development, dendrite morphogenesis, protein processing, and lipid transport (all p<0.001). On the other hand, discordant genes between human and mouse HH conditions in the NAc enriched for lysosome function, protein phosphorylation, transcriptional regulation, and ion transport (all p<0.001), while in the PFC these genes enriched for MAPK and ERK signaling, cell adhesion, and RNA processing (all p<0.001).

**Fig 8.**
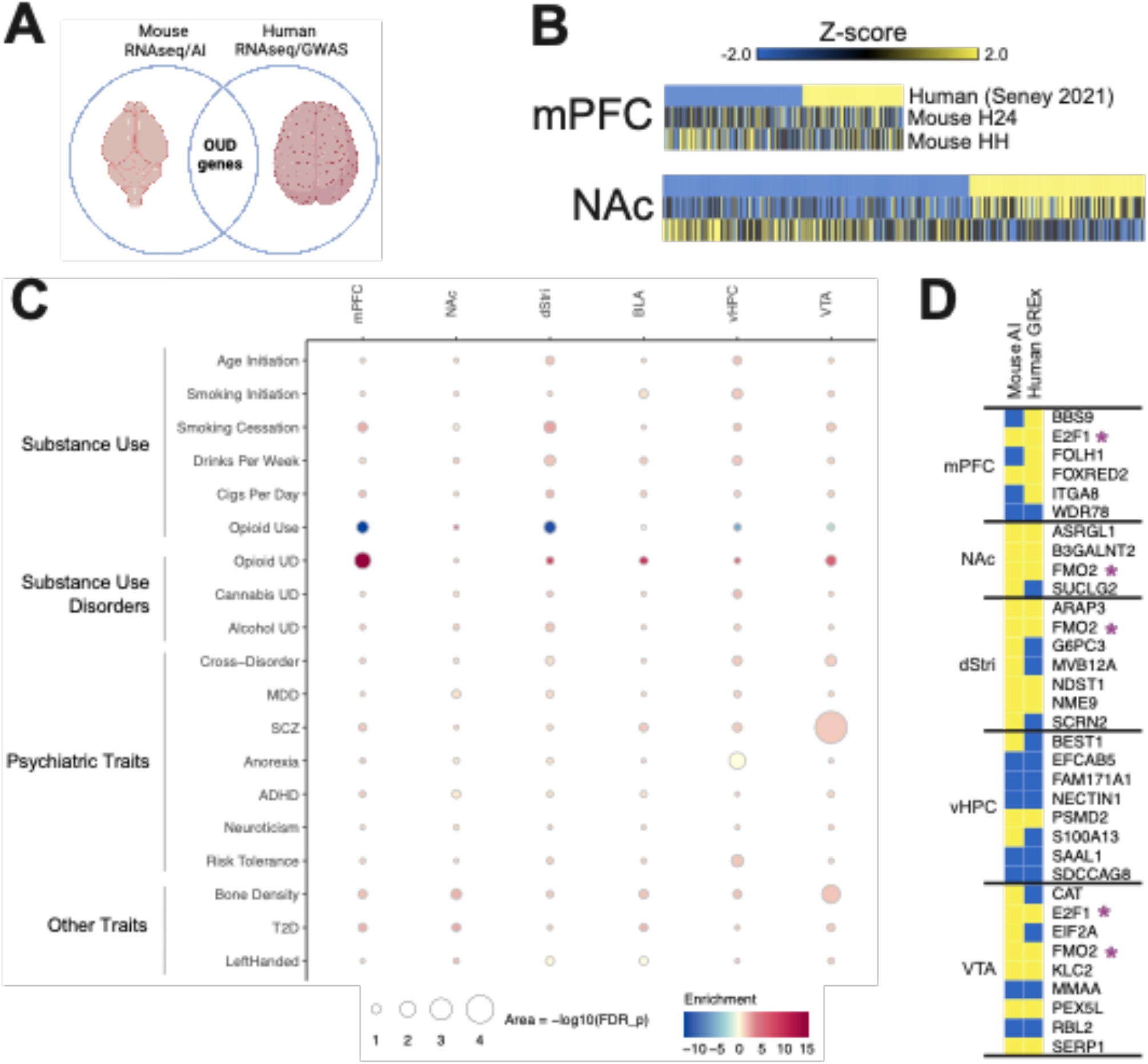
Convergent molecular drivers of OUD identified through integrated analysis of mouse and human genome-wide data. (**A**) Results from the present work were contrasted with available RNAseq data from postmortem tissue of humans with OUD and genome-wide association studies (GWAS) studies. (**B**) Union heatmaps showing gene expression relationships between human OUD patients (top row, downregulated in blue, upregulated in yellow; Seney 2021) and two relevant experimental conditions, H24 and HH, for mPFC and NAc. Gene lists were generated by combining common significantly regulated genes (z-score > |0.2|, p<0.05) between human and H24, and human and HH. (**C**) Association scores between AI-genes and GWAS traits related to Substance Use, Substance Use Disorders, Psychiatric Traits, and Other Traits. Columns show brain region, subdivided by negative or positive AI association. (**D**) Comparison of AI genes and tissue-specific GREx analysis. Overlap of genes identified as significantly (p<0.05) positively (yellow) or negatively (blue) associated with AI and GREx. Purple stars indicate genes that show overlap across multiple brain regions.

While genes identified in association with the AI reflect transcriptional correlates of an addiction-related behavioral phenotype in mice, GWASs identify genetic variants that associate with human behavioral phenotypes and disorder status at a population scale—results which can be leveraged to identify genomic regions associated with phenotypic risk. Since genetic variation predates lifetime exposures (e.g., drug, stress), GWASs may provide strong causal directionality in the development of addiction. Thus, integration of AI genes and GWAS hits could highlight key gene targets that serve as risk genes for addiction. We fist asked whether region-specific AI-associated genes contribute to risk for substance use or other psychiatric phenotypes using partitioned heritability linkage disequilibrium score regression (LDSC) (*15*). Genes positively and negatively associated with the AI in each brain region were tested for enrichment in SNP heritability for each trait using GWAS summary statistics (Fig. 8B). We observed strong enrichment for genes upregulated with increased AI in the mPFC and genes downregulated with increased AI in the VTA for OUD GWASs. Interestingly, we observed the opposite pattern of enrichment for opioid *use* GWASs. This suggests that AI-associated transcriptional signatures, especially those in mPFC and VTA, are likely to capture causal mechanisms of opioid OUD. Overall, AI gene enrichment was specific to OUD risk as opposed to other substance use and psychiatric disorders. This finding confirms the validity of using animal models of drug-intake to investigate mechanisms of addiction, and suggests that the molecular signatures captured in this study are primarily unique to OUD.

We then asked whether genes could be identified which are shared between AI and OUD risk. We translated OUD GWASs (OUD cases vs. healthy opioid-exposed controls; (*19*)) from the single nucleotide variant (SNV) level to the gene level using transcriptomic imputation. Specifically, we used S-PrediXcan (*18*) to impute brain-region-specific genetically-regulated gene expression (GREx) associations for OUD risk using GTEx models (*20*). These OUD GREx associations (OUD-GREx) reflect baseline transcriptional differences associated with a predisposition for OUD at the functionally translatable level of gene expression for comparison to our AI-associated genes. We found 31 genes that were significant (p<0.05) in both OUD-GREx and AI-associations with brain-region specificity—many of which were regulated in the same direction (Fig. 8C). Two genes, *FMO2* and *E2F1* were identified to be significantly positively associated with OUD-GREx and AI in multiple brain regions. *Fmo2*, flavin containing dimethylaniline monoxygenase 2, is involved in oxidative metabolism of xenobiotics which includes heroin (*32*). Little is known about *Fmo2* or its related gene partners in addiction, making it an interesting potential brain-wide therapeutic target to blunt drug intake. On the other hand, *E2F1*, E2F Transcription Factor 1, is part of a well-defined family of transcription factors involved in addiction-relevant behavior (*33–35*). Interestingly, *E2F1* has been implicated in coordinating transcriptional responses to fentanyl abstinence (*36*) and abstinence from morphine self-administration (*37*) within the NAc. Thus, these genes and others identified represent strong candidates as causal players in transcriptional mechanisms driving of OUD vulnerability.

## DISCUSSION

These studies combined heroin self-administration in mice, RNAseq in six brain reward regions, and advanced bioinformatic approaches to generate an atlas of transcriptional regulation relevant to OUD. We identified numerous transcriptomic changes across crucial reward processing centers (mPFC, NAc, dStri, BLA, vHPC, and VTA) driven by initial heroin exposure (SH), ongoing intake (H24), context-induced drug-seeking (HS), and relapse-like conditions (HH). This comprehensive, unbiased assessment of transcriptome-wide changes spanning multiple brain regions and exposure conditions enabled deep characterization of heroin-induced molecular dysfunctions that may promote drug-taking and prime relapse. Integration of addiction-relevant behavioral outcomes to create an AI with transcriptomic data revealed novel roles for brain regions, regulatory factors, and biological processes contributing to heroin intake. Further, comparisons of mouse RNAseq with that from OUD patients and GWAS results narrows in on key molecular targets with high therapeutic potential for OUD. These results serve as resource which can be leveraged for innumerable new avenues of research into molecular mechanisms and treatment of OUD.

Throughout these studies we noted that distinct brain regions play partly separable roles in coordinating molecular reprogramming by heroin exposure and context associations. Transcriptome-wide analysis using RRHO identified a strong coordination of gene expression between SH and HH conditions for the mPFC, NAc, and dStri, which was maintained at the level of DEGs. We also noted that SH and HH drive highly overlapping activation or inhibition of upstream regulator function in both mPFC and NAc, which was essentially absent for HS. The SH and HH conditions are similar in that heroin is on board prior to euthanasia. Coordination of transcriptional responses between these conditions implies a key role for these brain regions in orchestrating the effects of heroin. Additionally, genes primed across these two conditions may reflect key molecular mechanisms that undergo cyclical activation or suppression with repeated exposure to heroin and drive consolidation of drug-relevant motivational processes. On the other hand, we found that the BLA and vHPC exhibited a strong coordination of transcriptomic overlap between HS and HH, with particularly strong DEG overlap between conditions for vHPC. Interestingly, these regions did not show much upstream regulator engagement for HS, which could suggest that gene expression related to context-induced drug-seeking is more tightly controlled by fewer upstream regulators in comparison to conditions of relapse with heroin reexposure. Notably, AI-associated genes in the vHPC exhibit increasing engagement across H24, HS, and HH conditions indicative of a consolidation of gene priming with increasing exposure and relapse conditions. Taken together, these findings imply a separable compartmentalization of transcriptional regulation induced by heroin exposure itself vs. learned associations between heroin experience and exposure context. Cyclical activation of these gene networks may prime or desensitize genes through withdrawal periods, promoting vulnerability to relapse upon re-exposure to drug or drug-associated contexts.

Heroin altered numerous transcriptional regulatory patterns throughout the reward circuitry, but the molecular profile of the VTA stood out as unique. Transcriptome-wide analysis in the VTA identified no obvious bias for gene regulation induced by heroin exposure or context-induced drug-seeking. Compared to other brain regions, overall DEG expression in the VTA was moderate for both up- and down-regulated genes across experimental conditions. However, we found that acute heroin exposure (SH), ongoing intake (H24), context-induced drug-seeking (HS), and relapse-like conditions (HH) all caused a high degree of activation or inhibition of upstream regulator activity. Transcriptomic overlap from RRHO and pattern analysis found little bias between heroin exposure or context-induced seeking, but high degree of overlap in upstream regulators engaged by HS and HH. These results suggest that, although heroin does not cause striking DEG activation in the VTA, engagement of these DEGs may be controlled by multiple regulatory mechanisms. In contrast to the moderate level of DEG expression in the VTA, we found this region to have by far the most AI-associated genes. AI genes select for those that may have predictive power in promoting drug-taking and drug-seeking, suggesting that transcriptional changes within the VTA may directly promote ongoing drug-taking. This idea is consistent with findings integrating AI genes with human GWAS results which identified a strong relationship between AI genes and OUD. Further, genes positively associated with the AI were enriched for processes related to metabolic functions and energy utilization. These results are consistent with previous work using cocaine which identified distinct cocaine-induced transcriptional responses depending on exposure paradigm, but a convergent role for disruptions to energetic functions in mediating cocaine effects (*38*). These results suggest that molecular changes within the VTA may represent a pan-addiction target for disruption of drug intake and potentially drug-seeking.

One major theme uncovered by these studies was heroin-induced dysfunction in processes supporting ECM function throughout the reward circuit, which has been hypothesized to be a key component of OUD (*39*). The ECM of the brain is a complex milieu of proteins, proteoglycans, and polysaccharides that support various processes including cell-cell adhesion, blood-brain-barrier integrity, and synaptic plasticity (*30*). Here, we identified many genes and biological processes either directly or indirectly related to ECM remodeling, spanning collagen biology, chondroitin and keratan sulfate metabolism, and angiogenesis-related processes. Changes in collagen biology were particularly evident for conditions of ongoing heroin intake (H24). Interestingly, many genes that were found to be differentially regulated across multiple brain regions enriched for biological processes related to ECM functions. These changes were most notable in conditions of ongoing heroin intake and genes associated with drug-taking (AI-associated genes). Considering that the AI is heavily weighted to prior behavior (i.e., before withdrawal), these results may indicate that ECM changes are statedependent, and more specific to ongoing opioid use rather than driving relapse after protracted abstinence. These findings are consistent with previous reports that opioids induce changes to ECM biology (reviewed by(*39*)), specifically, recent reports from human OUD patients which identified a key role for heroin-induced changes to ECM biology in promoting opioid use (*14*). Thus, our findings confirm heroin-induced changes to ECM biology as a key driver of OUD, and extend this idea to cover largely the entire reward circuit.

The broad molecular reprogramming by heroin observed in these studies may be directly useful to inform existing knowledge of human OUD, and deconstruct relevant patterns of gene expression that promote drug-taking. One approach we took was to examine how our mouse RNAseq results directly related to recently developed human OUD RNAseq datasets from the mPFC and NAc. In the mPFC, we did not observe striking overlap between mouse and human conditions, but this may relate to any number of caveats, most obvious of which relates to anatomical ambiguity between rodent and human mPFC (*40*), and anatomical isolation of dorsolateral mPFC from humans versus infralimbic and prelimbic cortex from mice. On the other hand, we found strong overlap with the H24 condition in the NAc, but largely opposite regulation in comparison with humans in the HH condition. The ability of ongoing use to drive gene expression in one direction, but invert expression after re-exposure suggests that these genes undergo cyclical patterns of activation and inhibition through periods of abstinence and relapse that affect particular biological processes. Interestingly, we have recently noted a similar pattern of inversion of gene expression with re-exposure after abstinence when comparing RNAseq results from the NAc across cocaine IVSA in mice and postmortem tissue from humans with cocaine use disorder (*41*). Together, these studies solidify a crucial role for molecular reprogramming in the NAc to promote ongoing addiction-relevant behavior across drugs and species, and suggest that relevant gene expression networks undergo cyclical patterns of restructuring across abstinence and re-exposure conditions. Direct comparisons of gene regulatory networks engaged between periods of drug exposure and abstinence represents a crucial next step in identifying molecular drivers of addiction.

The AI developed in these studies enabled us to identify genes that may have predictive power in their ability to influence patterns of behavior relevant to addiction. This approach is similar to GWASs in humans which identify risk genes for addiction development. By integrating these two approaches, we identified positive enrichment of AI genes with OUD GWAS hits across all brain regions examined, indicating the utility of this approach in identifying addiction-relevant genes from rodent models. The strongest enrichment was observed for the mPFC and VTA, indicating that genes in these regions may be particularly susceptible to heroin-induced regulation to promote addiction. Interestingly, we noted that opioid use – i.e., not disordered use – exhibited mostly negative enrichment of AI genes with GWAS hits. These results could suggest that lower levels of these AI genes may have a protective effect against the development of problematic opioid consumption indicative of OUD. Notably, enrichment of AI genes was largely specific to opioid conditions, further emphasizing that this dimensionality reduction approach effectively narrowed analysis to opioid-relevant processes. To identify predictive OUD candidate genes throughout the reward circuitry with this AI approach, we used transcriptomic imputation to convert OUD GWASs to gene-level variants which revealed brain-region-specific GREx associations for OUD risk. This approach revealed 31 genes that were significantly different in both AI and GREx across 5 brain regions, many of which were regulated in parallel. Interestingly, two genes, *FMO2* and *E2F1*, were identified to be affected in multiple brain regions, and were consistently positively associated with addiction-relevant outcomes in both AI and GREx across all regions. E2F family transcription factors have recently been implicated in cocaine addiction in rodent models, further highlighting their likely importance in addiction mechanisms (*24, 33, 34*). Thus, these genes may hold exceptionally high promise at potential novel therapeutic targets for treating OUD.

In conclusion, the present studies identified numerous genes, regulatory systems, and biological processes affected by heroin exposure, intake, seeking, and relapse. Throughout the brain reward circuitry, we uncover networked and compartmentalized transcriptional changes that may drive drug-taking and drug-seeking. Dimensionality reduction approaches to integrate transcriptomic findings and behavioral outcomes refined key addiction-relevant gene expression signatures that promote overlapping functional changes across brain reward regions. Further, we show that these data are directly applicable to the human condition and can be leveraged to isolate convergent targets relevant to OUD. These results present a broad landscape of heroin-induced molecular reprogramming relevant to OUD that can be mined to uncover new mechanisms of addiction and potentially inform novel treatment strategies.

## ETHICS DECLARATION

The authors declare no competing financial interests

## ACKNOWLEDGEMENTS

This work was supported by funding from the National Science and Research Council of Canada (Postdoctoral Fellowship to CJB) and the National Institute of Health (P01DA047233 to EJN).

---

**Fig S1.**
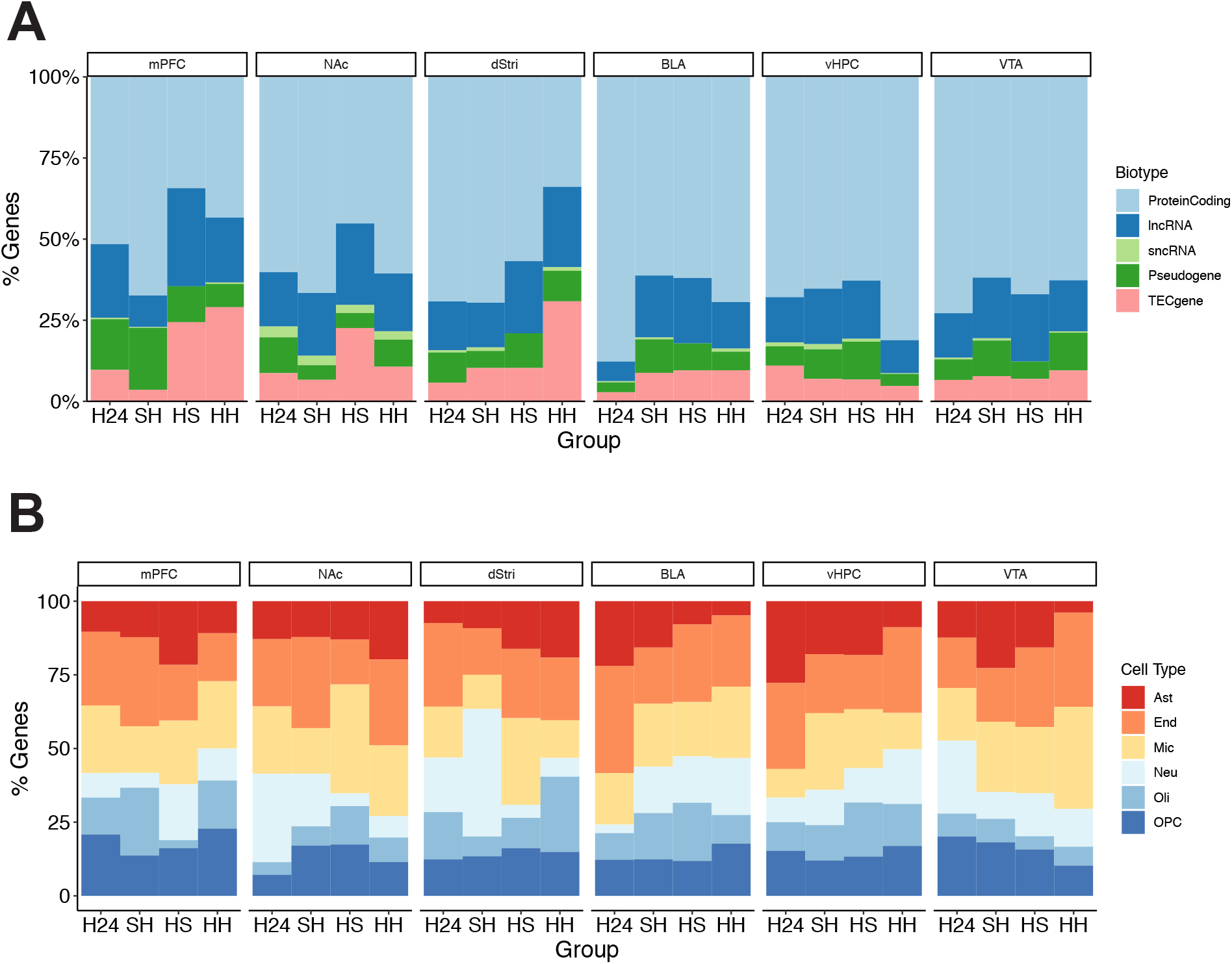
Heroin self-administration causes region-specific transcriptional regulation across gene biotypes and cell types throughout the brain’s reward circuitry. (**A**) Proportion of transcripts belonging to particular biotypes (protein coding, long noncoding RNA (lncRNA), short noncoding RNA (sncRNA), pseudogene, or to be experimentally confirmed (TEC)) presented across experimental conditions faceted by brain region. (**B**) Proportion of differentially expressed genes enriched for cell-type-specific markers of astrocytes (Ast), endothelial cells (End), microglia (Mic), neurons (Neu), oligodendrocytes (Oli), or oligodendrocyte precursor cells (OPC) presented across experimental conditions faceted by brain region.

**Fig S2.**
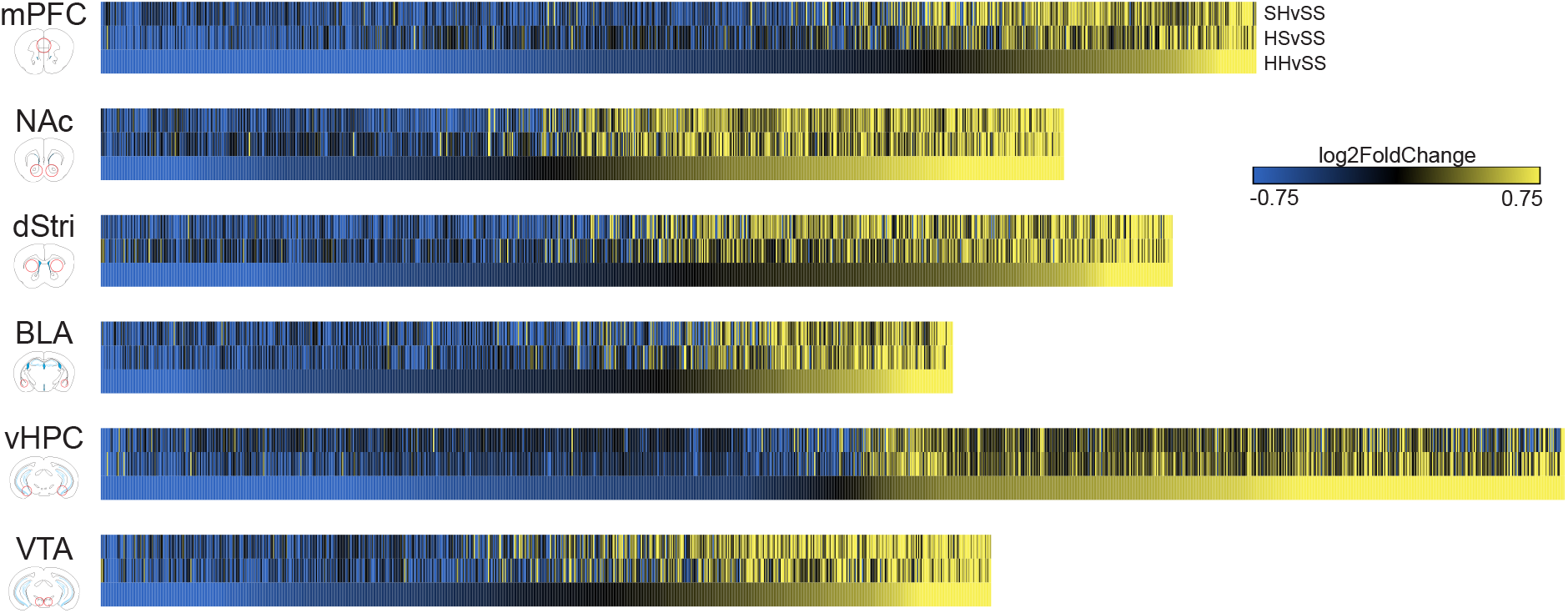
Differential gene expression induced by heroin IVSA after prolonged abstinence. Union heatmaps seeded to log2FoldChange of the HH condition (heroin IVSA followed by 30 d withdrawal and an acute heroin challenge) showing broadly similar transcriptional regulation for up- or downregulated genes across the SH condition (acute heroin exposure: saline IVSA, 30 d withdrawal and an acute heroin challenge) and the HS condition (heroin withdrawal: heroin SA followed by 30 d withdrawal and an acute saline challenge) across all brain regions.

**Fig S3.**
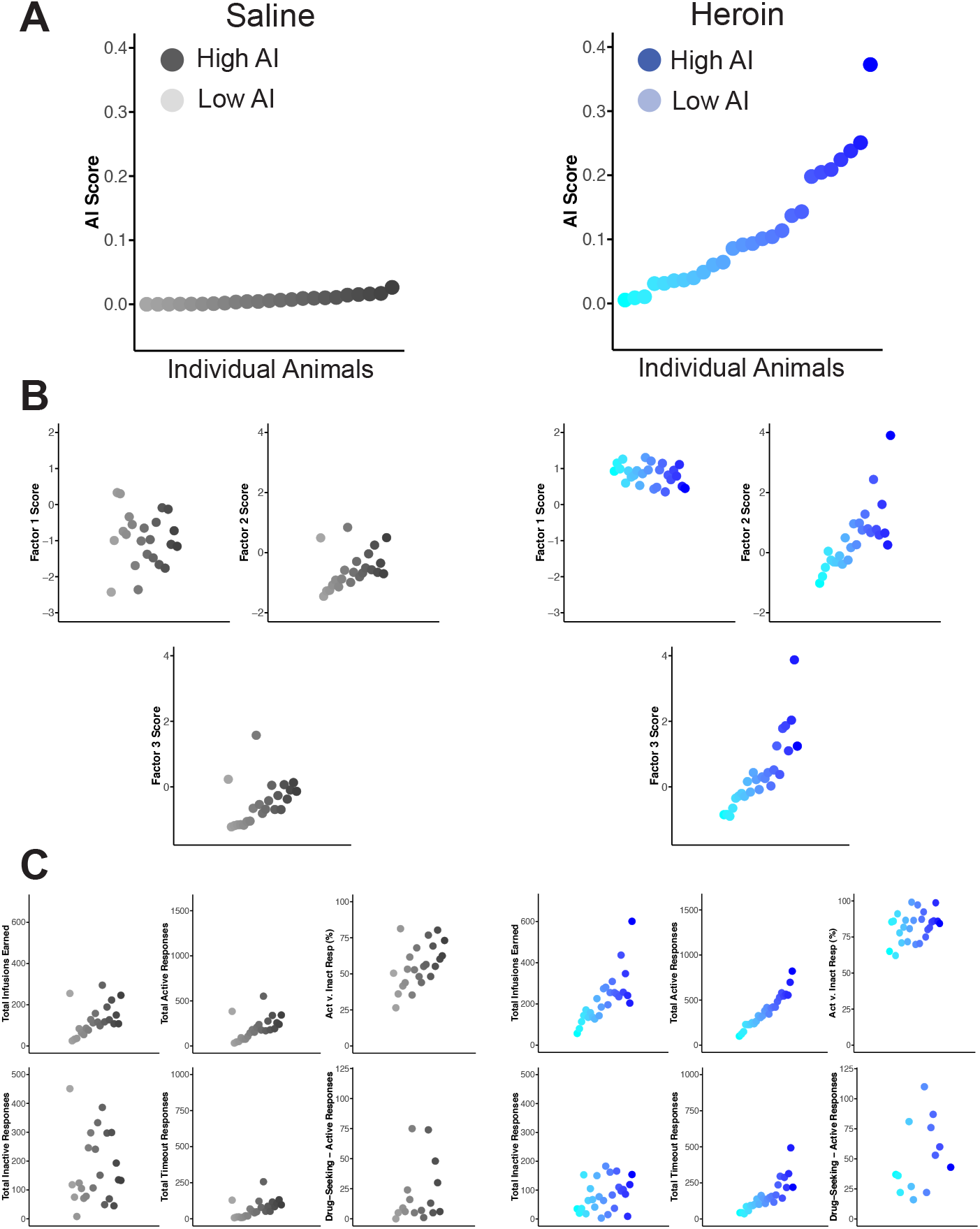
The addiction index (AI) effectively separates animals with and without a history of heroin intake across behavioral domains and latent variables identified through Factor Analysis. (**A**) Plots show the AI score attributed to individual animals represented as gray (saline IVSA; left) or blue (heroin IVSA; right) dots, staggered for visibility. Saturation of symbol color represents relative AI score in each group, and is carried through the following figure panels. (**B**) Plots show individual animals colored by AI score (saturation, as in panel A) in either saline (gray; left) or heroin (blue; right) mapped onto values for individual Factors identified, showing that higher AI (more saturation of symbol) is associated with higher Factor. Note that Factor scores are generally higher for heroin animals compared to saline animals. (**C**) Plots show individual animals colored by AI score (saturation, as in panel A) in either saline (gray; left) or heroin (blue; right) mapped onto individual behavioral measures used for Factor Analysis. Note that with the exception of Total Inactive Responses heroin animals show higher scores for all behavioral measures, and animals with higher AI tend to score higher on each behavioral variable.

**Fig S4.**
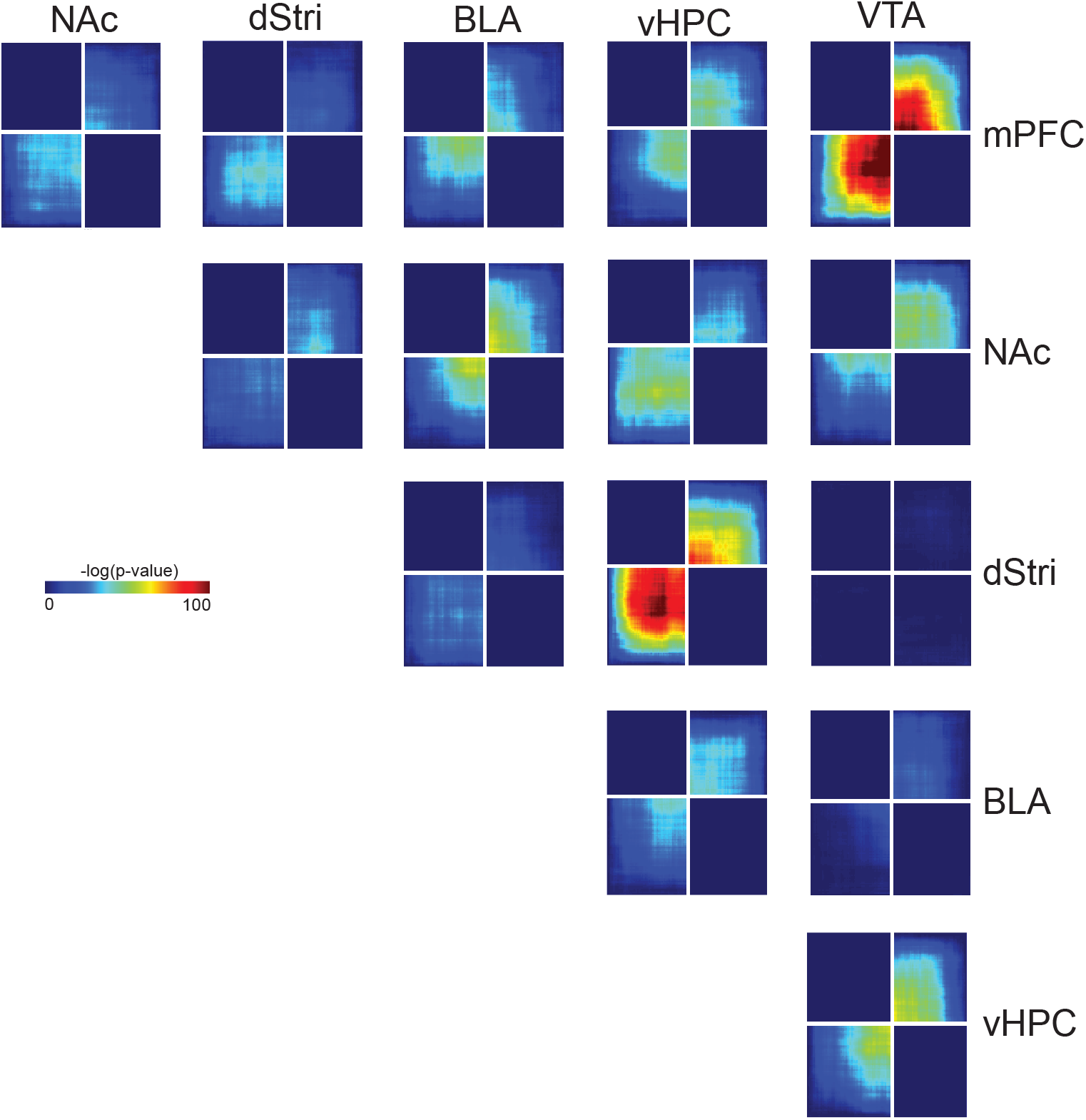
Genes identified to be associated with the Addiction Index are largely independent across brain regions. Rank-rank hypergeometric overlap plots comparing all brain regions to one another reveal generally independent AI-associated gene expression patterns, but identify a strong concordance between the VTA and mPFC, and the vHPC and dStri.

